# Rewarding, but poorly rewarded: Gendered narratives of science communication in the life sciences

**DOI:** 10.1101/2024.02.28.582614

**Authors:** Perry G. Beasley-Hall, Pam Papadelos, Anne Hewitt, Charlotte R. Lassaline, Kate D. L. Umbers, Michelle T. Guzik

## Abstract

Science communication (*sci-comm*) encompasses activities that promote scientific literacy, inform policy, and inspire future scientists. Despite its value, sci-comm is often considered less pretigious than research and other internal academic activities. In Australia, scientific organisations (e.g., learned societies) rely on members to perform a range of sci-comm activities, typically unpaid. In this pilot study, we surveyed 88 Australian life sciences organisations to examine who performs sci-comm and why. Most respondents were early-career women employed in university research positions. Participants largely agreed that organisational sci-comm brought limited career benefits and was often viewed as feminised or “care work” relative to their research roles. Yet, respondents also cited personal and professional gains and most wished to continue in such roles. Our findings suggest “invisible” sci-comm performed for scientific organisations is disproportionately undertaken by women at early career stages, with implications for career progression and gender equity in STEM.

## 1 Introduction

Science communication (hereafter *sci-comm*) is broadly defined as work that seeks to increase scientific literacy and raise public awareness of science by making research findings accessible to a wide range of audiences (Lewenstein, 2015; van Stekelenburg *et al*., 2022). Science communicators perform diverse activities which have only been grouped under a single descriptor relatively recently (Davis, 2014; Gascoigne & Metcalfe, 2017). For this reason, definitions in the scientific literature vary. Sci-comm is often used as shorthand to refer to activities performed by academics to a non-scientific audience, also called *outreach* (Davis, 2014; Johnson *et al*., 2014; Woitowich *et al*., 2022), but may also include activities performed by individuals in sectors outside of academia, such as government agencies, museums, NGOs, or science centres (Gascoigne & Metcalfe, 2017; Delicado, 2020), or directed at a scientific audience rather than the non-scientist public (also called *inreach*; Bik *et al*., 2015). For consistency with the broader scientific literature, here we will use the term to refer to outreach performed by scientists primarily employed in university settings.

Sci-comm can positively influence science policy (Hajdu & Simoneau, 2020), shape the career choices of aspiring scientists, and improve retention rates of university students (Peckham *et al*., 2007; Wrighting *et al*., 2021). As such, some scientists view sci-comm as an ethical responsibility integral to their day-to-day work (Martín-Sempere *et al*., 2008). The skills gained by practicing sci-comm may also be considered as an asset for career progression or networking (Daoust-Boisvert, 2022). However, broader attitudes towards sci-comm are not always positive, and scientists may not be adequately rewarded for communicating their research. Scientists working in academia often hold ambivalent or negative attitudes towards sci-comm due to a perception that time spent on outreach detracts from the quality of scientific research, impeding future promotion success or career progression more generally, a stigma termed the “Sagan effect” (Johnson *et al*., 2014; Watermeyer, 2015; AbiGhannam, 2016; Negretti *et al*., 2022). Sci-comm is also not often included in university workload models and instead typically performed on an unpaid basis as a “labour of love” (Therkildsen, 2022; Docter-Loeb, 2024; Schøning *et al*., 2025). An additional challenge facing science communicators is the persistence of gendered narratives in sci-comm as a field. Women are systematically underrecognised in science in favour of men (Knobloch-Westerwick *et al*., 2013) and sci-comm is generally considered a woman-dominated field (Johnson *et al*., 2014; Delicado, 2020). This has led to it being typified as a “soft”, feminised, and caring practice distinct from the production of “expert” pedagogical or research material typically associated with men (Johnson *et al*., 2014; Pérez-Bustos, 2014). Further, women are more likely to experience gendered stereotypes and harassment when communicating science compared to men (Amarasekara & Grant, 2019; McKinnon & O’Connell, 2020).

Given the diverse attitudes and benefits associated with sci-comm, a knowledge gap remains regarding which demographics engage in this work, what motivates them to do so, and the associated benefits given its typically unpaid nature. One mode of sci-comm understudied in the literature is that performed on behalf of scientific organisations, which we here define to include learned societies, professional associations, non-profits, or clubs. These bodies are usually run by, and cater to, university academics. As such, we consider them an exemplar of behaviour and attitudes in academia more broadly, particularly because they have a clearly defined service dynamic. Sci-comm activities performed for scientific organisations are diverse and may include, but are not limited to, giving public talks, managing a social media or website presence, planning events (e.g., networking events and competitions), media production, and administrative duties. It is our anecdotal experience that most people who engage in sci-comm in this context do so on a voluntary basis as members of the organisation in question; this is consistent with science communicators typically receiving little to no financial compensation for such work, which is done in their spare time (Riedlinger *et al*., 2019; Negretti *et al*., 2022). We consider these roles “invisible” labour, a term first coined to refer to women’s unpaid work in the home, but more recently extended to service tasks in academia (Sliwa *et al*., 2021; Gordon *et al*., 2024). In this context, invisible sci-comm can be considered labour contributed to an organisation that is unseen, as is its impact (Social Sciences Feminist Network Research Interest Group, 2017).

Australia has a unique history in its support of science communication and was among the first in the world to offer postgraduate courses in the field (Gascoigne & Metcalfe, 2017). The 1990s saw the formation of Australian Science Communicators (ASC), the first ever representative body for the field globally, as well as initiatives such as Science Meets Parliament and Questacon’s Science Circus (Gascoigne & Metcalfe, 2017). The establishment of the ASC was spurred by a lack of data surrounding funds and time spent on sci-comm in Australia. Yet, few studies have examined demographics who engage in sci-comm in Australia and why (McKinnon & O’Connell, 2020), let alone “invisibly” for organisations. This is particularly relevant given sci-comm is increasingly performed on online platforms where authorship, time investment, and impact can be difficult to measure or attribute (Duffy & Sawey, 2022). The present study began with a dual aim: to examine who undertakes sci-comm work in Australia and why, and to examine an assumption held by the authors that invisible roles performed on behalf of organisations are typically held by early-career researchers (ECRs), many of whom are women. We therefore sought to explore the following in a pilot study: 1) which demographic(s) undertake science communication work in scientific organisations in Australia; 2) the responsibilities and motivations associated with such roles; and 3) the perceived value of such work for career progression. To investigate these questions, a survey was disseminated to Australian scientific organisations and clubs associated with the life sciences— the study of microorganisms, plants, fungi, and animals— asking respondents about their experiences, motivations, and perceptions of sci-comm work carried out on behalf of such organisations. We supplement our findings with select insights from a companion study that interviewed a subset of participants surveyed here (Papadelos & Beasley, 2025).

## 2 Methods

From December 2022 to March 2023, we distributed a self-administered online survey to life sciences societies in Australia. To identify relevant scientific organisations, we used a keyword-based web scraping approach. We conducted a systematic Google search using the keywords “society” or “organis(z)ation” alongside life sciences disciplines including biology, zoology, botany, physiology, evolution, ecology, entomology, ornithology, ichthyology, mammalogy, genetics, and microbiology. The first two pages of Google results were visited and reviewed to produce a list of 88 Australian life sciences organisations. Contact information for the relevant president or chair of each organisation was collected to facilitate a request that they distribute the survey to their members.

The survey was hosted online using the Survey Monkey platform and consisted of 11 questions that were either closed, multiple choice or open-ended questions. Data was collected on respondents’ demographic background, including how they would characterise their career stage (early, mid, or late-career). The open-ended questions sought information about the nature of sci-comm contributions to life sciences organisations including respondents’ perceptions of the value of such work. Because of the small sample size of the target population, no formal sampling strategy was used. Instead, scientific organisations were encouraged to disseminate the survey by email to third parties using snowball sampling to maximise the number of respondents (Patton, 1990). We called for responses from members of these organisations irrespective of their involvement in sci-comm initiatives in the past, with the intention of gauging perceptions from both those with direct sci-comm experience and individuals involved in life sciences organisations more broadly. The survey was followed by six in-depth interviews with people who were currently, or had recently, carried out sci-comm work for a scientific organisation, the results of which are presented in Papadelos and Beasley (2025). Together, this approach combined quantitative survey data with qualitative information from the interviews. After the data had been analysed separately, triangulation was used “to obtain different but complementary data on the same topic” (Creswell and Clark, 2007, p. 62). Our understanding of the perspectives and experiences of those involved in sci-comm in scientific organisations was enhanced by this method. An examination of the survey data is detailed below.

## 3 Results

### 3.1 Who undertakes sci-comm work for life sciences organisations?

A total of 49 responses were received through the online survey. Of the 49 respondents, 28 identified as women (57%), 20 as men (41%), and one as a non-binary person (2%). In relation to career type, 20 respondents (40%) described themselves as university academics. The remainder comprised non-academic scientists working in sectors such as the public service (26%), non-university academics employed in institutions such as museums (16%), those with non-science careers (4%), and students (4%). Among those who stated that the question of career stage was applicable, the majority—25 respondents (64%)—identified as early- or mid-career researchers (EMCRs), while the remaining 14 participants (36%) described themselves as late-career researchers (Fig. 1).

**Figure 1.**
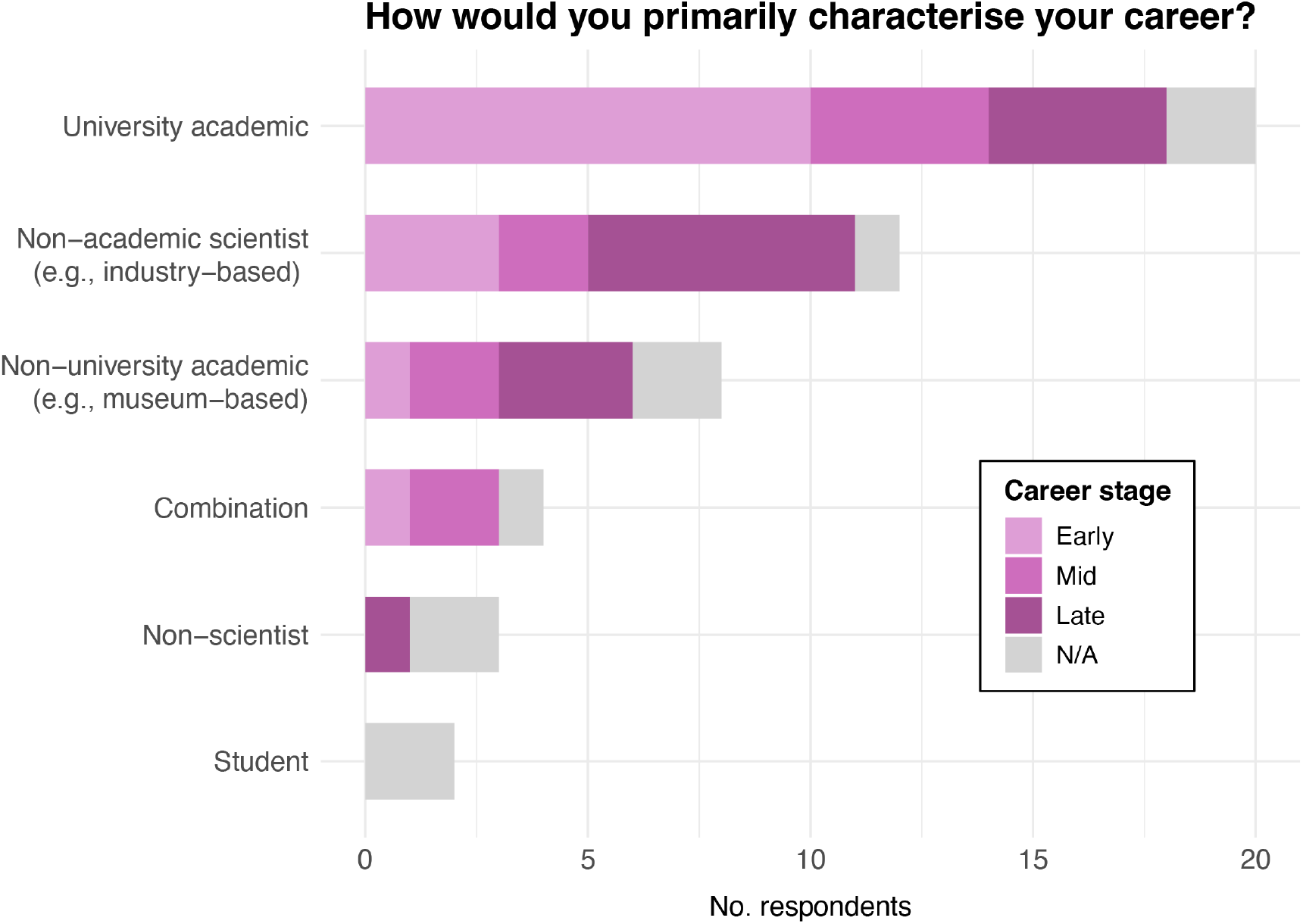
Career type and stage of respondents who participated in the online survey. The “combination” category refers to respondents who identified with more than one career type.

Taking these responses in combination, the majority of survey participants were women early or mid-career researchers (EMCRs) employed at an Australian university. Of the 49 respondents, 44 (89.8%) reported having previously engaged in sci-comm activities on behalf of a scientific organisation. The organisations corresponded to a diverse set of fields within the life sciences, with the most reported fields being entomology, ecology, and biology (Figure 2). Several respondents reporting multiple disciplines as their primary field. Additionally, most respondents indicated that the scientific organisation to which they contributed sci-comm work was associated with more than one scientific discipline.

**Figure 2.**
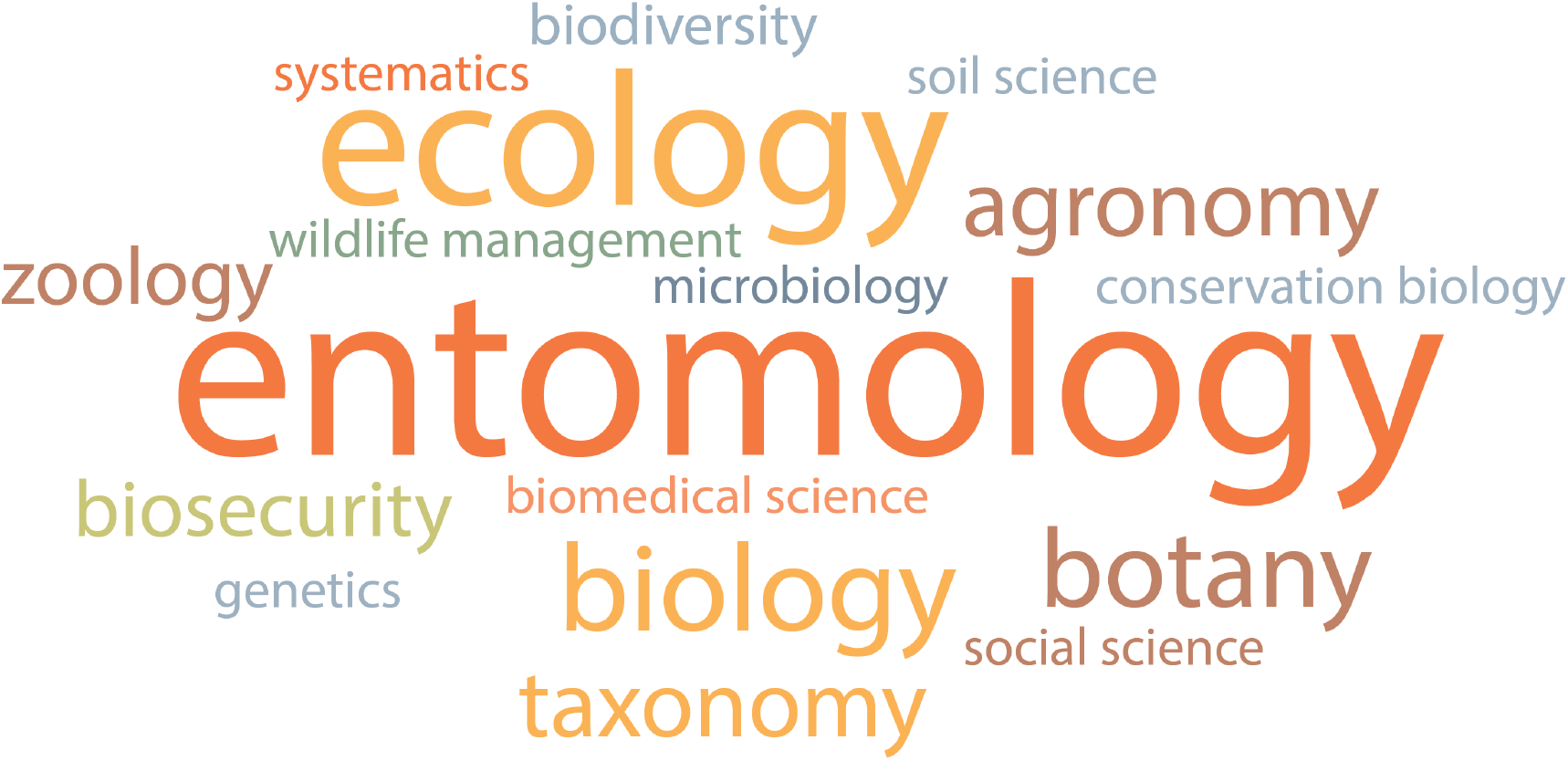
Word cloud of scientific fields of organisations for which respondents had performed sci-comm activities.

### 3.2 The nature of sci-comm contributions

Media production, such as the production of newsletters or posters, was the most common type of sci-comm work performed on behalf of life sciences organisations and was reported by 54.5% of respondents, over half of which were women. The next most common contribution was public speaking and social media engagement followed by administrative tasks, advertising, event organisation, and journal editing. In terms of gender distribution, women were most prominently represented in social media management (72.8%), public speaking (61.5%), and media production (58%). Administrative tasks were generally shared between men and women, while the only respondent who had contributed as a journal editor was a man. Excluding journal editing, two women and one man reported contributing to all of the sci-comm activities listed above.

Most respondents reported performing sci-comm work on an unpaid basis (81.8%). Among those who had contributed to paid roles, over half were men (62.5%). All respondents who had undertaken paid sci-comm roles reported that their contributions were acknowledged through authorship in scientific publications. Men and women were equally represented among the 71% of respondents who reported that they worked on sci-comm both during and after regular business hours. Of the eight respondents who said they did this work exclusively outside of work hours, five were women and one identified as non-binary. Notably, only women reported performing this work exclusively during their working hours. Four respondents (three women and one man) did not respond to this question. Furthermore, over half of the respondents (59.2%) reported receiving no mentorship or assistance in such roles.

### 3.3 Perceptions and motivations surrounding sci-comm work

The extent of acknowledgement and perceived value of sci-comm work were markedly different within and outside of the scientific organisations concerned. Of the 44 respondents who had performed sci-comm work for organisations, 37 (84.1%) reported that their contributions were acknowledged in some way by the societies, including via journal articles, newsletters, annual reports, or websites. Several participants stated that instead of formal recognition they were verbally acknowledged or received gifts for their contributions. In contrast, just over half (23, 52.2%) of respondents stated that their contributions to life sciences organisations had not been acknowledged in an academic context. The minority of respondents who did receive academic recognition reported that this ranged from their contributions being recognised during promotion interviews to being awarded research prizes.

Most respondents did not view their contributions to be important in advancing an academic career (Figure 3). A Likert scale was used to assess the value of sci-comm work within an academic context, with responses ranging from “not at all” (26.5%), “a little” (24.5%) or “somewhat” (22.5%) “very” (14.3%), or “extremely” (4.1%). Almost all respondents who rated sci-comm “not at all” valuable for academic career progression were women (84.6%), while most who answered “a little” or “somewhat” were men.

**Figure 3.**
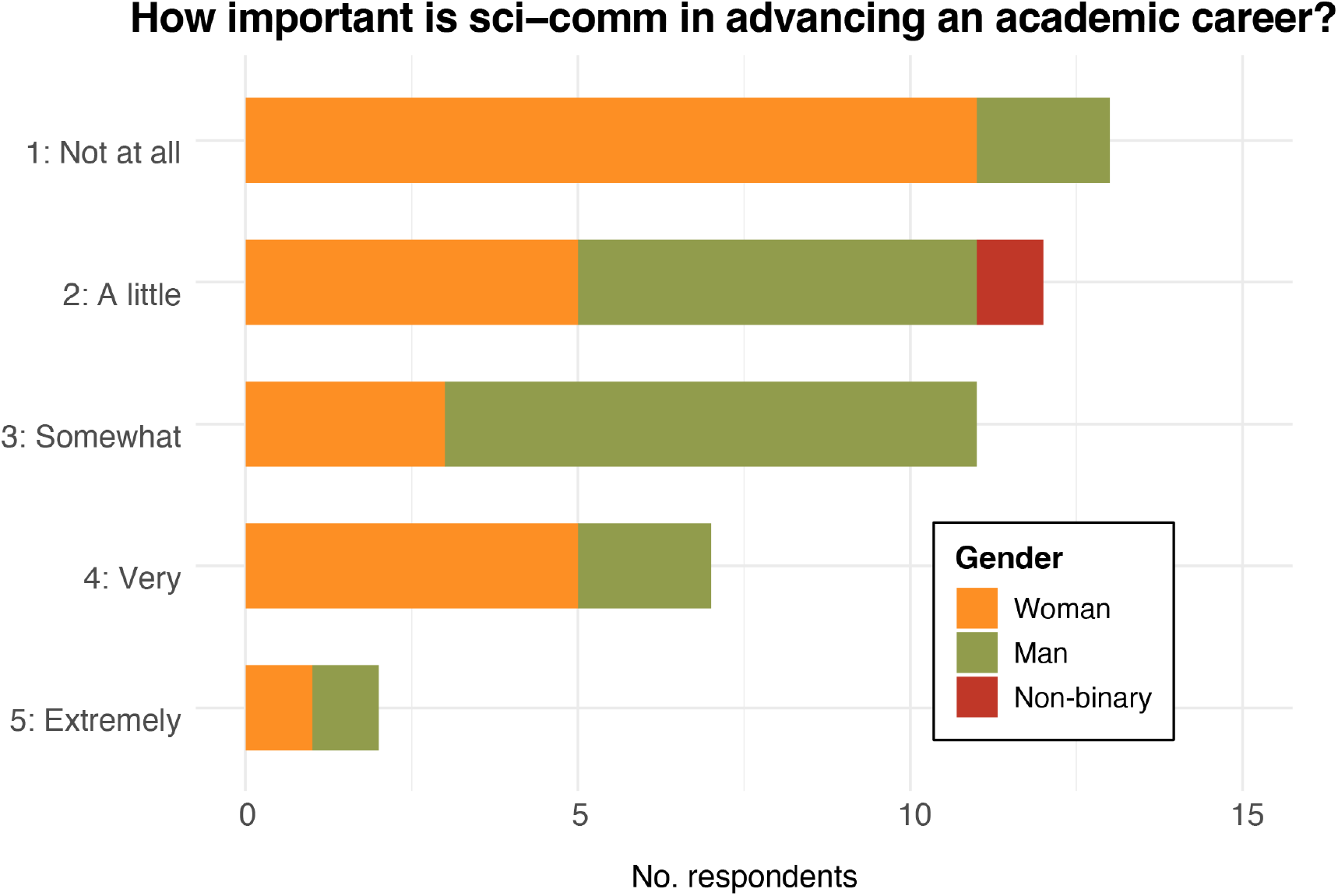
Attitudes of respondents towards the value of sci-comm activities for academic career advancement.

Overall, while most respondents expressed satisfaction with the impact of their sci-comm work, many conveyed doubts about its value in advancing their academic careers. Among those who identified advantages to engaging in such work, the most frequently cited motivations were skill development and personal fulfilment (43%). This was followed by networking opportunities (20%), a desire to explain and promote science to the general public (20%), career progression (11%), and a sense of obligation to give back in recognition of the opportunities and benefits gained through participation in the scientific community (4%; these values do not total 100% as some respondents reported more than one benefit). One respondent claimed that sci-comm work held no value for them, describing it as merely part of a “busy period on top of usual responsibilities” (Woman, state government scientist, MCR). Several respondents offered further insight into how the lack of formal recognition for sci-comm within academic settings had negatively impacted them. For example:

> *Early career people might go into these opportunities for their career advancement. You soon realise that, whilst extremely valuable experience, it is not acknowledged enough by those who pay your salary*. (Man, university-based researcher, ECR)
>
> *Industry, including government, value these skills highly - but they were not valued in academia*. […] *It’s almost a self-fulfilling prophecy - do this ‘extra’ work to make yourself more employable outside of academia just in case, but by doing so you are likely to make yourself less competitive in academia*. (Woman, university-based researcher, ECR)
>
> *Once you have developed this skill set and are not paid or recognised for your time, however, it begins feeling much less valuable and quite exploitative*. (Man, university-based researcher, ECR)
>
> *Exploitation is rife in his industry but we, as science communicators, often have no choice but to engage with this if we would like to develop professional relationships and realise our motivations for entering the field (be it for career purposes or altruism)*. (Man, university-based researcher, ECR)

Almost all respondents stated they would continue with sci-comm contributions if they were given the opportunity to do so (87.2%), despite the “non-trivial time commitment” (Man, university-based researcher, career stage not stated). Respondents articulated clear drivers for doing sci-comm work beyond direct career benefits, such as skill development, gaining experience and confidence in public speaking, and making new social connections for future collaborations:

> *The value I receive is in my own education. I have no formal training in this field so every article I add to the society’s newsletter is a learning opportunity*. (Woman, non-scientist, retired)
>
> *It’s helped me to make connections outside of my immediate network and in different fields*. (Man, university-based researcher, ECR)

Others were motivated by personal satisfaction derived from communicating research, promoting the importance of science and evidence-based decision-making, and educating the public. Some respondents noted that it was fulfilling regardless of (limited) career benefits, and it provided them satisfaction to contribute to the broader image of science in society:

> *I receive personal satisfaction. I strongly believe that being a scientist is a tremendous privilege, and part of our responsibility is to ensure our work is communicated back to our communities. I also believe I have a responsibility to be visible as a role model for women and minority communities*. (Woman, university-based researcher, MCR)
>
> *I find it fulfilling to see how this work impacts others. The exposure to these topics and the people that study them can be limited, yet it is important to make it available to others. You have no idea what positive impact it may have*. (Woman, university-based researcher, ECR)
>
> *I mostly do it for the community and the public …I’m not convinced academia values or recognises sci comm yet. I can’t be sure that the inclusion of my sci comm work in my successful promotion application did make a difference, but I am sure it helped a bit to present me as a well-rounded scientist*. (Woman, university-based researcher, MCR)

Finally, some respondents saw clear professional benefits in performing sci-comm work for scientific organisations, such as being recognised in promotions, awards, or grant applications in their paid roles:

> *Because of the research work and the publication, I was able to obtain a full scholarship to follow my research passion*. (Woman, university-based researcher, ECR)
>
> *As an academic, there was some professional value in that it could be used for promotion, award and grant applications and yearly work plans*. (Man, university-based researcher, MCR)
>
> *I think it is well-received by job and promotions committees. Its* [sic] *just another type of expected service*. (Woman, non-university academic, MCR)

The results above indicate a disconnect between the personal and professional value of sci-comm work. While many respondents—particularly women EMCRs—were actively involved in a range of sci-comm activities and reported personal satisfaction and skill development as key benefits, this work was often unpaid, unrecognised, and perceived as having limited impact on academic career advancement.

## 4 Discussion

Here, we conducted a pilot study to better understand the demographic(s) engaging in sci-comm activities on behalf of life sciences organisations in Australia, particularly with respect to gender, career stage, and career type. We were interested in the nature of these roles, why individuals performed in them in the first place, and the perceived value of this type of invisible sci-comm in academic and non-academic contexts. Although sci-comm is generally recognised as the professional responsibility of scientists (Martín-Sempere *et al*., 2008; AbiGhannam, 2015; Schøning *et al*., 2025), our data revealed persistent inequities in who undertakes this work and how it is acknowledged and rewarded. Of the 49 respondents surveyed in the present study, most were women early to mid-career researchers working in university settings. Over half of respondents (59%) reported that they received no training to perform in a sci-comm role on behalf of a scientific organisation and the majority (82%) did so unpaid.

### 4.1 “Invisible” sci-comm is gendered

Our preliminary findings demonstrate that gender and career stage play a major role in who performs sci-comm on behalf of scientific organisations in Australia. Most respondents worked in university settings, where there is extensive literature showing that women are disproportionately assigned teaching and administrative duties (Misra *et al*., 2011; Guarino & Borden, 2017). Within this context, sci-comm is often categorised as administrative work that is commonly ignored. A noticeable gender dynamic emerged in our data, with women more likely to take on invisible sci-comm tasks—such as social media management or internal communications—while men occupied public-facing, prestigious roles, such as journal editing. This division reflects broader gendered patterns prevalent throughout academia in which women undertake communal, service-oriented roles and men occupy individualised, public-facing positions (Johnson *et al*., 2014; Wilkinson *et al*., 2022). These disparities are rooted in social norms that position women as primarily responsible for care and service work, both at home and in the workplace (Wilkinson *et al*., 2022). As a result, even highly educated women are often guided towards lower-status, poorly compensated roles, particularly when that work is collaborative, supportive, and carried out for the benefit of the organisation rather than for individual recognition (Beasley & Papadelos, 2024).

Intersecting with the gendered narrative discussed above is our finding that invisible sci-comm roles are disproportionately performed by early to mid-career researchers. This is perhaps unsurprising as women are overrepresented at early career stages in academia and are typically employed on short-term, precarious contracts, whereas fixed-term and permanent researchers at senior levels are predominantly men (Social Sciences Feminist Network Research Interest Group, 2017; Coin, 2018; Rosser *et al*., 2019). These results are consistent with previous work demonstrating junior researchers are more likely to undertake invisible sci-comm than their more senior colleagues (Martín-Sempere *et al*., 2008; Riedlinger *et al*., 2019; Dudo *et al*., 2021). We note that other studies have found engagement in sci-comm instead increases with seniority (Jensen *et al*., 2008; Bentley & Kyvik, 2011; Jensen, 2011). However, this discrepancy may be due to much of the existing literature examining sci-comm undertaken on an individual basis, which is more likely to be performed by senior staff, rather than on behalf of scientific organisations as discussed here. Indeed, limited research on the intersection between career stage and sci-comm activity supports this idea: senior academics are more likely to engage in “one-time-only” opportunities of higher prestige (promoting their own research), whereas junior academics are more likely to engage in education outreach in schools (disseminating the research of others) (Jensen *et al*., 2008). Ultimately, while sci-comm is viewed as essential, there is a clear gendered dimension to how scientists engage with this work. This dynamic has the potential to perpetuate discrimination, exploitation, or marginalisation of those who undertake such roles, particularly along gendered lines.

### 4.2 Sci-comm is rewarding, but poorly rewarded

Respondents expressed mixed views regarding their reasons for undertaking sci-comm on behalf of scientific organisations. Two major motivations for engaging in sci-comm work emerged in our survey results. The first was associated with a sense of responsibility to one’s field in the life

sciences or society as a whole. Respondents reported that they were motivated by assisting those at more junior career stages, translating scientific concepts to the general public, being visible as a role model to minorities in science, and promoting their field of science. As one PhD student (woman) remarked, any personal inconvenience is acceptable as “the value to society is priceless.” The second motivation was personal and included the development of transferable skills (such as communicating research), networking, and attracting students. Other personal motivations included self-growth, “warm fuzzies” from interacting with the non-scientist public, and the enjoyment of speaking to an enthusiastic audience. These narratives are consistent with previous studies demonstrating motivations for engaging in sci-comm may be personal or driven by a sense of duty as a scientist (Martín-Sempere *et al*., 2008; Cerrato *et al*., 2018; Daoust-Boisvert, 2022; Woitowich *et al*., 2022; Schøning *et al*., 2025). Respondents derived a high degree of satisfaction from sci-comm activities performed for organisations, with 84% indicating a desire to continue in such roles in future.

The benefits of engaging in sci-comm discussed above exist in tension with a lack of broader institutional recognition. While most respondents received recognition from the organisation for which they had performed sci-comm work, this was typically not the case for in their day-to-day roles, particularly those in university settings. Respondents working in universities frequently compared sci-comm to other feminised academic roles like teaching and administration (Doerr, 2022), reflecting broader patterns around societal expectations of unpaid labour (Alhammad, 2022) This type of invisible work has been described as “academic housekeeping” and is rarely, if ever, associated with career advancement (Jensen *et al*., 2008; Ecklund *et al*., 2012; AbiGhannam, 2015; Lidskog & Standring, 2024). In general, respondents did not consider their sci-comm contributions to be valuable to academic career progression, and women generally held a more negative view of the value of this work than men (Figure 3). Similar sentiments were expressed by those not working in universities. One non-academic scientist described having to “ghost-write for organisation promotion” (man, late career). Others pointed to systemic limitations that prevented meaningful acknowledgement of individual contributions: after an organisation encouraged its members to engage in sci-comm as volunteers, one respondent declined to participate because the body was not prepared to waive the cost of membership in exchange for her labour (woman, student).

### 4.3 How can sci-comm engagement be made more equitable?

The introduction of policies intended to redress gender inequalities in higher education is a constructive step toward institutional reform and has been encouraged by organisations such as Science in Australia Gender Equity (SAGE), the Athena Scientific Women’s Academic Network (SWAN), and ADVANCE (Rosser *et al*., 2019; McKinnon, 2022). Nevertheless, these initiatives have yet to produce meaningful or lasting change. This is clearly reflected in the ongoing underrepresentation of women in senior leadership positions within higher education, despite decades of activism from feminist academics (Barclay *et al*., 2025). Scholars have devoted considerable attention to understanding why this is the case. According to O’Connor (2023), an important contributing factor is “legitimating discourses”, a term describing the normalisation of existing (inequitable) policies and practice as fair and reasonable that, in turn, obscures gendered power at a structural level. Examples include that women inherently lack excellence, leadership, or confidence to facilitate access to higher positions; that women simply choose not to prioritise career advancement; or that organisations are inherently fair and unbiased. These narratives clearly rely on biological essentialist narratives and ignore gendered power imbalances on a structural level. By shifting the responsibility to redress gender inequalities away from institutions, such discourses instead present gender gaps as the natural consequence of personal decisions or inherent characteristics, thereby placing the pressure on individuals to improve by making better choices (O’Connor, 2023).

Insights from a companion qualitative study reinforce this link between legitimating discourses and discrimination (Papadelos & Beasley, 2025). As stated previously, a subset of respondents was interviewed to gain a more nuanced understanding of this issue. There was a consensus among the six participants (five women and one man) that sci-comm work for scientific societies was “invisible”, “thankless”, and “constant.” Although most found it emotionally rewarding, they also emphasised the personal cost of taking on this invisible labour—particularly the time lost to research or other career-advancing tasks. Nonetheless, the women interviewed (*n* = 5) unanimously agreed that this work should not be mandated or included in workload expectations, as requiring scientists to do so might compromise its quality. They suggested that, although formal recognition could be beneficial, the primary focus should be on addressing underlying structural issues. According to one interviewee, the problem is not just a lack of visibility but the fact that the benefits of sci-comm are widely shared, while the burdens are unequally distributed. Papadelos and Beasley (2025) therefore argue that sci-comm is a form of institutional care mainly provided by women. This is a complex issue, as women have historically assumed caregiving roles across both private and public spheres and many value the relational dimension of building connections with disparate audiences. However, its frequent invisibility, lack of compensation, and time-intensive nature can result in long-term resentment and contribute to systemic discrimination (Beasley & Papadelos, 2024). In order to mitigate this problem, the authors recommend a series of structural reforms in academia which include the creation of formal job sci-comm titles, integration of sci-comm into hiring and promotion processes, and the inclusion of care and relational labour within workload models (Papadelos & Beasley, 2025). Further recognition of sci-comm work could also be encouraged by scientific organisations themselves, namely proper attribution of individual contributions and the provision of financial compensation where appropriate and possible (Papadelos & Beasley, 2025).

### 4.4 Limitations of this study

While the current study offers important insights into the experiences of scientists engaged in invisible sci-comm work, several challenges and limitations must be acknowledged. Our findings are situated within the Australian context, where sci-comm contributions on behalf of scientific societies are often voluntary and unpaid, and this may differ in international settings. We also focused on organisations related to the life sciences as a case study to explore broader academic attitudes, but we recognise that scientists working in these subfields, especially biology, are among the most active science communicators (McCullagh *et al*., 2019). As such, the gender gap in sci-comm participation we observed here may not be directly comparable to other fields, such as the physical sciences (Ecklund *et al*., 2012), and limit the applicability of our findings. Moreover, there is a possibility that not all survey respondents shared a consistent understanding of the term *science communication*, particularly given diverse definitions of the term in the scientific literature. This was evident in instances in which respondents mentioned being recognised in journal publications or acting as a journal editor, which would typically apply to inreach rather than outreach, the focus of the present study. Finally, we note that the sample size and nature of our survey may have influenced our results, particularly as individuals who do not engage in sci-comm frequently may have perceived the study as less relevant to them and may have been less likely to participate (self-selection bias). Our results were limited to gendered narratives affecting women as only one non-binary person participated in the study. Overall, our preliminary findings demonstrate the need for further research exploring these questions with a larger sample size, including gender diverse individuals, and representing broader academic disciplines and organisations.

## 5 Conclusions

While this study surveyed individuals working in non-academic and academic settings in Australia, academic contexts were of particular interest to us as institutions have developed a broad range of frameworks to encourage gender equity, yet women in academia continue to experience barriers to career progression. One such example is the undervaluing of sci-comm work, even though it is acknowledged as being crucial to the translation of complex research. Sentiments around sci-comm work remain poorly understood or quantified, and questions remain about who takes on these roles and why, particularly for academics given their apparent negative connotations. To address these concerns, and to take up recent calls for more research on the value of contemporary sci-comm (Autzen & Weitkamp, 2019; Wilkinson *et al*., 2023), we investigated motivations associated with sci-comm undertaken on behalf of Australian life sciences organisations (which tends to be volunteer work done by researchers) and the perceptions attached to such work in an academic context.

This study supports past research demonstrating that women may be disproportionately penalised when taking on voluntary roles, such as sci-comm, due to perceptions that they reduce research capacity or indicate less academic rigour (particularly when performing in a service or “invisible” role). Our preliminary findings suggest most sci-comm roles are performed by early to mid-career women scientists working in universities. The majority of respondents expressed that sci-comm work is important and essential and was usually acknowledged by the science organisation involved. In contrast, participants consistently expressed that it was undervalued or inadequately acknowledged in academia. Most respondents, regardless of gender, were aware that sci-comm work was not factored into their workload and did not support their career progression, with some indicating that it might be detrimental. Nonetheless, the majority indicated that they would continue doing such work because of a range of associated personal and professional benefits. For this reason, we argue that it is important to address inequities in the way that sci-comm work is recognised and rewarded.

## 7 Acknowledgements

The authors have no conflict of interest to disclose. We would like to thank Chris Beasley for providing helpful suggestions on an earlier version of this manuscript and Natalie Thomas in her role as a research assistant. This study was funded by a University of Adelaide Faculty of Arts, Law, Business, and Economics interdisciplinary research grant and received ethics approval from The University of Adelaide Human Research Ethics Committee (no. H-2022-175).

